# Effects of transmembrane phenylalanine residues on γ-secretase-mediated Notch-1 proteolysis

**DOI:** 10.1101/2024.12.11.628002

**Authors:** Shweta R. Malvankar, Michael S. Wolfe

**Affiliations:** Department of Medicinal Chemistry, School of Pharmacy, The University of Kansas, Lawrence, KS 66045

**Keywords:** Intramembrane proteolysis, sequence specificity, immunoblotting, mass spectrometry

## Abstract

γ-Secretase is a presenilin-containing intramembrane aspartyl protease complex that cleaves within the transmembrane domain (TMD) of nearly 150 substrates, with the amyloid precursor protein (APP) being the most well studied. APP cleavage by γ-secretase generates amyloid β-peptides (Aβ) that pathologically deposit in Alzheimer’s disease. APP TMD substrate undergoes initial endoproteolysis (ε-cleavage) followed by processive carboxypeptidase trimming of long Aβ intermediates in ∼tripeptide intervals. Although γ-secretase cleavage of Notch1 is essential in developmental biology and altered in many cancers, the processing of this cell-surface receptor is relatively understudied. Only one sequence specificity rule is known for γ-secretase substrate processing: Aromatic residues such as phenylalanine are not tolerated in the P2’ position with respect to any processing event on the APP TMD. Here we show using biochemical and mass spectrometry (MS) techniques that this specificity rule holds for Notch1 as well. Analysis of products from the reactions of purified enzyme complex and Notch1 TMD substrate variants revealed that P2’ Phe relative to ε-site cleavage reduced proteolysis and shifted initial cleavage N-terminally by one residue. Double Phe mutation near the ε site resulted in reduced proteolysis with shifting to two major initial cleavage sites, one N-terminally and one C-terminally, both of which avoid Phe in the P2’ position. Additionally, three natural Phe residues were mutated to corresponding residues in the APP TMD, which led to increased ε proteolysis. Thus, Phe residues can affect the enzyme reaction rate as well as cleavage site specificity in the Notch1 TMD.

## INTRODUCTION

γ-Secretase is an intramembrane-cleaving protease which hydrolyzes the transmembrane domain (TMD) of its substrates within the hydrophobic environment of the lipid bilayer.^1^ The protease is a complex of four evolutionarily conserved integral membrane proteins: nicastrin, APH-1, Pen-2 and presenilin, the catalytic component of the complex.^2,3,4^ The human γ-secretase complex is now known to have over 145 substrates,^5^ including the amyloid precursor protein (APP) and Notch1 receptor. Dominant missense mutations in APP or presenilins, particularly presenilin-1 (PSEN1) cause early-onset familial Alzheimer’s disease (FAD),^6^ implicating altered processing of APP substrate by γ-secretase to pathologically deposited amyloid β-peptide (Aβ) in triggering the disease process.^7^ For this reason, the proteolytic processing of APP substrate by γ-secretase has been extensively studied. APP ectodomain shedding by β-secretase produces the 99-residue membrane-bound C-terminal fragment C99, which serves as the substrate for γ-secretase in the production of Aβ peptides. C99 undergoes initial endoproteolytic cleavage by γ-secretase at the ε-site, producing 49- or 48-residue long Aβ intermediates Aβ49 or Aβ48.^8^ These peptides then undergo processive trimming by γ-secretase, generally in tripeptide intervals, producing Aβ peptides along two pathways: Aβ49→Aβ46→Aβ43→Aβ40 and Aβ48→Aβ45→Aβ42→Aβ38.^9^ Much less is known about γ-secretase processing of Notch1, arguably the most biologically important of all substrates of the protease complex.

The Notch family of cell-surface receptors is evolutionarily conserved and essential to all metazoans, being involved in cell proliferation, cell fate determination, physiological homeostasis and damage repair.^10,11,12,13,14^ Dysregulated Notch signaling leads to tumorigenesis as well as progression of various diseases. ^12,13,15,16, 17^, γ-Secretase-mediated TMD cleavage is a critical step in Notch signaling (Fig. 1). Initial cleavage at the S1 juxtamembrane site in the secretory pathway leads to a heterodimeric receptor that traffics to the plasma membrane. Interaction with cognate ligand on a neighboring cell triggers proteolysis of the heterodimeric receptor at the S2 site by the metalloprotease ADAM10, resulting in ectodomain shedding^18,19,20^ and leaving behind the Notch Extracellular Truncation (NEXT) in the membrane.^21^ NEXT then undergoes cleavage by γ-secretase at the TMD S3 site, near the cytosolic interface and comparable to ε cleavage in APP C99,^22^ to release the Notch intracellular domain (NICD)(Fig. 1).^23,24^ The NICD then translocates to the nucleus, where it initiates gene expression.^25^ The remaining membrane-bound stub of Notch, a fragment dubbed Nβ that is comparable to Aβ49, is further cleaved by γ-secretase at S4 sites to release shorter Nβ peptides.^26^ Presumably, long Nβ is processively trimmed to short Nβ in a manner similar to Aβ49 trimming to Aβ40, but this has not been formally demonstrated.

**Figure 1:**
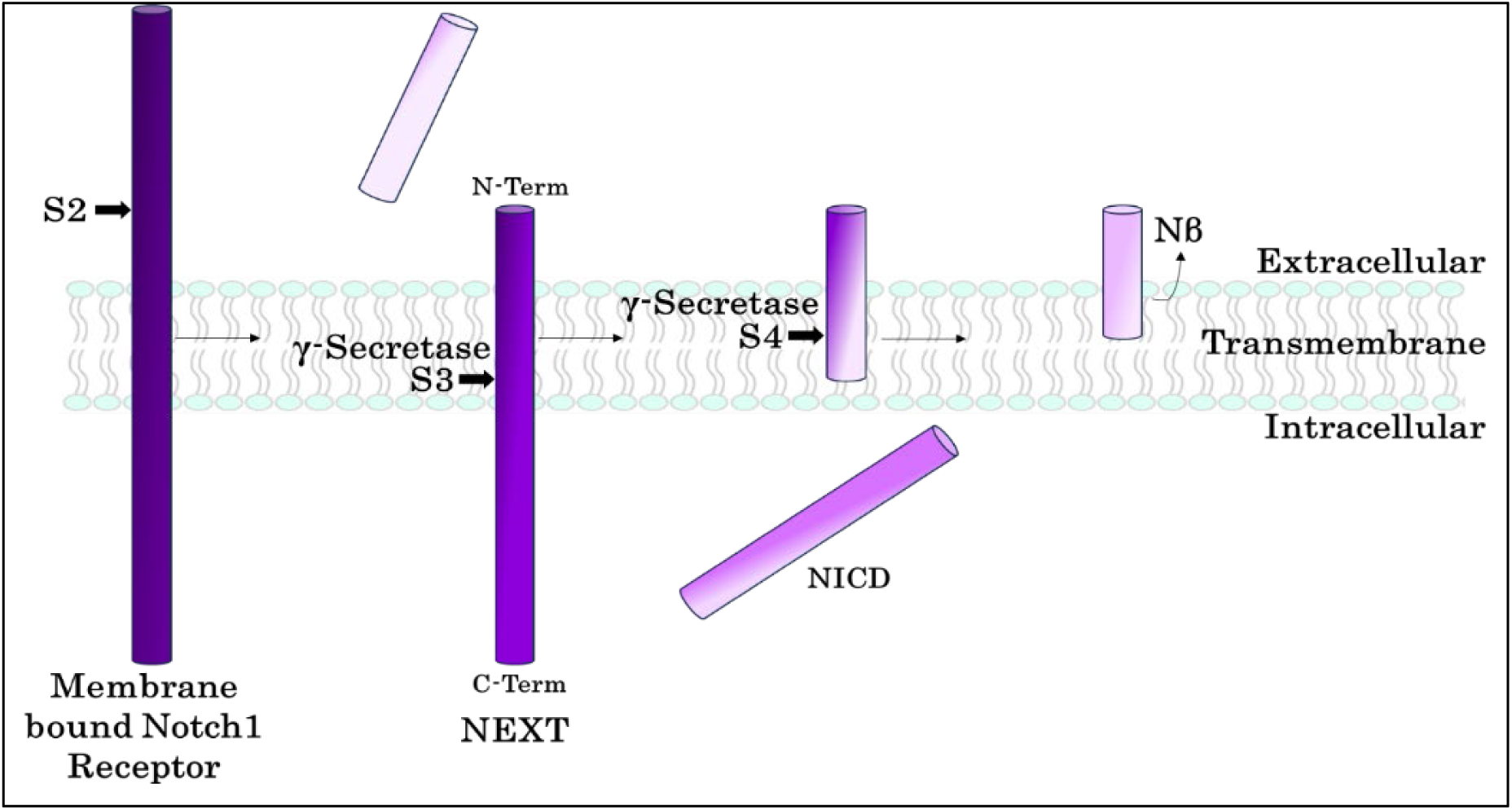
Proteolysis of Notch1 by γ-secretase. After the S2 cleavage of Notch1 by ADAM10, the remaining membrane-bound fragment of Notch1, called <u>N</u>otch <u>Ex</u>tracellular <u>T</u>runcation (NEXT), undergoes first cleavage by γ-secretase at the S3 site in the TMD near the cytosolic interface. This cleavage in turn releases the <u>N</u>otch Intra<u>c</u>ellular <u>D</u>omain (NICD) from the membrane. The remaining fragment undergoes further cleavage by γ-secretase at S4 sites to release Nβ products extracellularly, the Notch counterpart of Aβ peptides produced from γ-secretase processing of APP substrate.

No consensus sequence in substrate TMDs has been described for γ-secretase.^27,28,29^ The only sequence specificity rule known is that bulky aromatic residues, particularly Phe, are not tolerated in the P2’ position with respect to any cleavage site within the APP TMD, apparently due to a shallow corresponding S2’ pocket in the protease active site.^30,31^ However, whether this “phenylalanine rule” applies to any of the other γ-secretase substrate is not known. Here we tested this specificity rule with Notch1, combining mass spectrometry and biochemical analyses. We found that γ-secretase does not tolerate Phe in the P2’ position relative to the S3 (ε) cleavage site in the Notch1 TMD, reducing cleavage in addition to shifting the initial cleavage site. We also discovered that the natural Phe residues in the Notch TMD sequence are important for efficient S3 cleavage by γ-secretase. Furthermore, we observed that the final S4 cleavage sites were not altered for any of Notch1 variants under investigation. These findings are nevertheless consistent with tripeptide trimming coupled with the P2’ Phe specificity rule.

## RESULTS

### Design and γ-secretase processing of wild-type Notch1 (Notch_WT) substrate

We and others previously showed that the TMD of Notch1 substrate is sufficient for processing by γ-secretase and that proteolysis is slowed by extending the extracellular domain.^27,32^ We therefore designed a recombinant substrate based on the human Notch1 sequence composed of the TMD flanked on either end by short juxtamembrane regions (Fig. 2A), similar to a previously used Notch1 sequence for NMR studies^33,34^. A 6XHis tag was added to the N-terminus, and a FLAG epitope was added to the C-terminus, to facilitate isolation and detection of N- and C-terminal cleavage products. The N-terminal juxtamembrane Notch1 sequence is the same as previously reported for structural studies of Notch-bound γ-secretase and begins at the natural S2 cleavage site that generates NEXT.^21,35^ The schematic diagram also depicts the expected NICD- and Nβ-like products that would be generated by γ-secretase. All recombinant NEXT-based proteins in this study include a translation-initiating N-terminal methionine, previously reported to have little or no effect on binding and cleavage by γ-secretase.^35^

**Figure 2.**
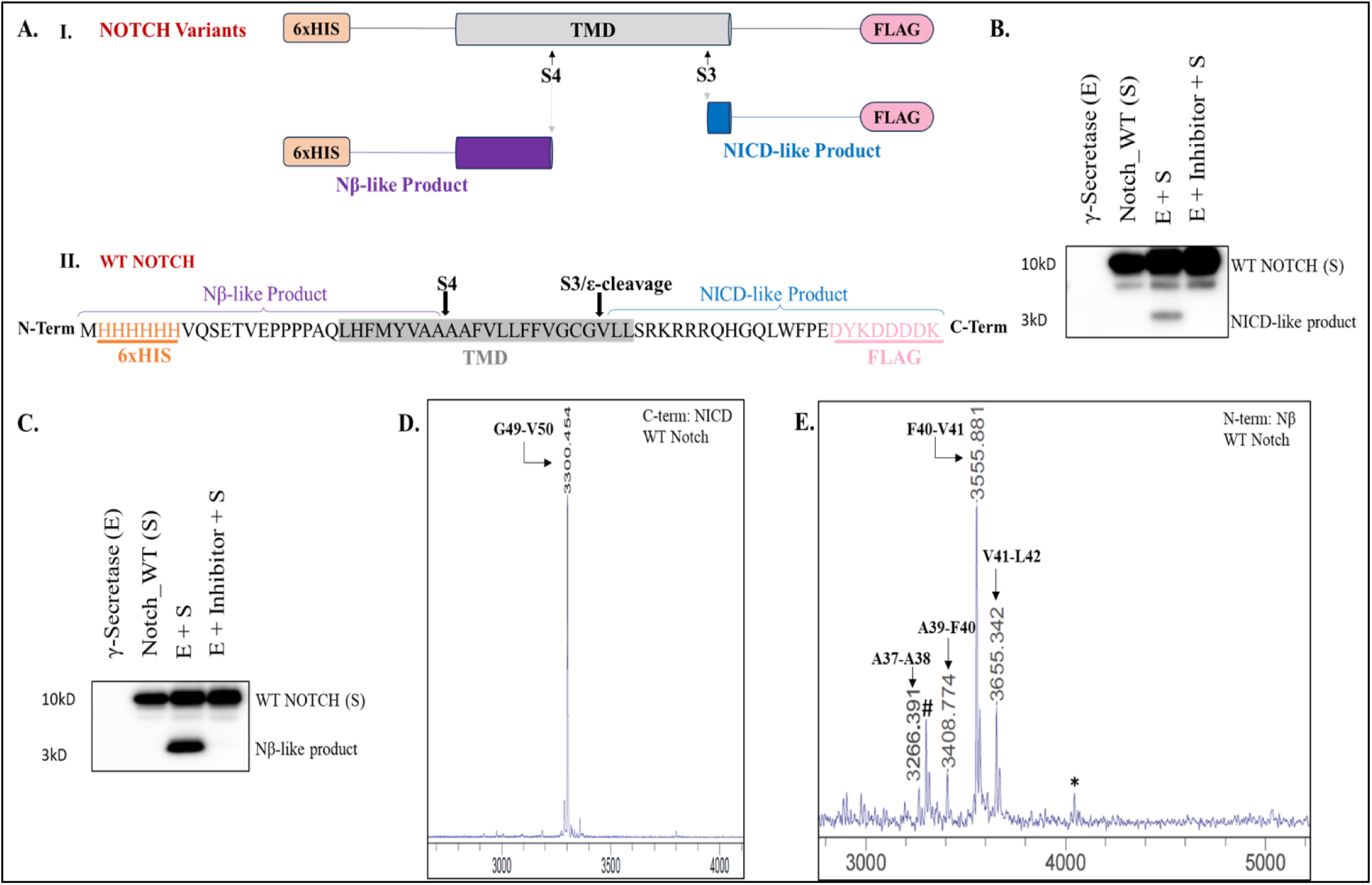
WT Notch1 substrate and analysis of products of γ-secretase processing. A. Schematic representation of truncated and tagged WT Notch1 substrate and expected NICD-like and Nβ-like products. Also shown is the complete WT Notch1 substrate sequence used in this study, with the reported S3 and S4 cleavage sites by γ-secretase. B. Immunoblot analysis of γ-secretase processing of WT Notch1, using anti-Flag antibody to detect NICD-like products. C. Immunoblot analysis of γ-secretase processing of WT Notch1, using anti-His antibody to detect Nβ-like products. In both B and C, no product formation was observed in presence of γ-secretase inhibitor (I) or with enzyme or substrate alone. D. MALDI-TOF MS spectrum of immunoprecipitated NICD-like C-terminal product from incubation of WT Notch1 substrate with enzyme. Calculated mass: 3299.691, observed mass: 3300.454. E. MALDI-TOF MS spectrum of immunoprecipitated Nβ-like N-terminal products from incubation of WT Notch1 substrate with enzyme. Calculated mass for major product species: 3552.67205, observed mass: 3555.881. *Represents unidentified peak and # represents a C-terminal product non-specifically bound and eluted from beads. Other labels indicate the scissile amide bond leading to the specific product.

As proof that the designed NEXT-based sequence is processed normally by γ-secretase and suitable for further study, we first expressed the recombinant wild-type protein (WT Notch) from *E. coli* BL21 cells and purified the protein by affinity chromatography through the Flag epitope tag. The identity and purity of the substrate was confirmed by SDS-PAGE and matrix-assisted laser desorption/ionization time-of-flight mass spectrometry (MALDI-TOF MS)(Fig. S1) and then subjected to proteolytic reaction with recombinant WT γ-secretase enzyme complex expressed and purified from suspension human embryonic kidney (HEK) 293 cells as previously described.^31,36,37^ After incubating 30 nM enzyme with 10 µM WT Notch at 37 ^0^C for 4 h, NICD- and Nβ-like product formation was confirmed by western blotting using anti-Flag antibody (for NICD-like product; Fig 2B) and anti-His antibody (for the Nβ-like product; Fig 2C). NICD-like and Nβ-like products were only observed in the enzyme plus substrate reaction mixtures, and product formation was blocked by γ-secretase inhibitor LY411,575,^38,39^ indicating that product formation was dependent solely on enzyme action.

To confirm if both the NICD- and Nβ-like products indeed result from cleavages occurring at the known S3 and S4 cleavage sites of Notch1, we analyzed these products by MALDI-TOF MS. The enzyme reaction was subjected to immunoprecipitation using anti-FLAG antibody for analyzing NICD-like products and anti-His antibody for analyzing Nβ-like products. The MALDI-TOF spectrum of NICD-like product from enzyme incubated with WT Notch substrate clearly showed a single cleavage product at the known S3 site between amino acids G1753 and V1754 (human full-length Notch1 numbering)(Fig. 2D).

Based on the first cleavage site by γ-secretase for human Notch1 and APP, we aligned the TMDs of human Notch and APP as shown in Table 1. We assumed that this first cleavage site (S3) of human Notch by γ-secretase is analogous to the major Aβ49 ε-cleavage, making the Notch S3 cleavage between G49 and V50. All the Notch sequence numberings in this study are assigned based on the alignment with the APP C99 sequence (Table 1). The MALDI-TOF spectrum of Nβ-like products immunoprecipitated from the same enzyme substrate reaction showed cleavage at the previously reported major S4 site between A37 and A38 (Fig. 2E).^26^ In addition to this known cleavage, we also observed Nβ-like peaks corresponding to other cleavage sites such as A39-F40, F40-V41and V41-L42, as shown in Fig. 2E and Table 2, and are remarkably similar to the cleavage sites reported previously in cellular studies.^26,40^ However, in our case, the Nβ-like peak from cleavage between F40-V41 is more intense than the peak from A37-A38 cleavage. Based on our numbering from Table 1, F40-V41 cleavage is analogous to cleavage of APP C99 leading to production of Aβ40, the major Aβ variant of γ-secretase processing. Also, we did not observe any peaks corresponding to Nβ product generated directly from S3 cleavage (Fig. 2E and Table S1).

**Table 1.** Alignment of sequences in and around the TMD of human Notch1 and amyloid precursor protein (APP). Sequences were aligned based on the first cleavage by γ-secretase, between residues 49-50 at the ε site for APP and at the S3 site for Notch1. Numbering is according to APP C99 γ-secretase substrate that produces Aβ peptides.

| Numbering | 29 | 30 | 31 | 32 | 33 | 34 | 35 | 36 | 37 | 38 | 39 | 40 | 41 | 42 | 43 | 44 | 45 | 46 | 47 | 48 | 49 | 50 | 51 | 52 |
| --- | --- | --- | --- | --- | --- | --- | --- | --- | --- | --- | --- | --- | --- | --- | --- | --- | --- | --- | --- | --- | --- | --- | --- | --- |
| APP | G | A | I | I | G | L | M | V | G | G | V | V | I | A | T | V | I | V | I | T | L | V | M | L |
| Notch 1 | Q | L | H | F | M | Y | V | A | A | A | A | F | V | L | L | F | F | V | G | C | G | V | L | L |

S3/ ε-cleavage

**Table 2.**
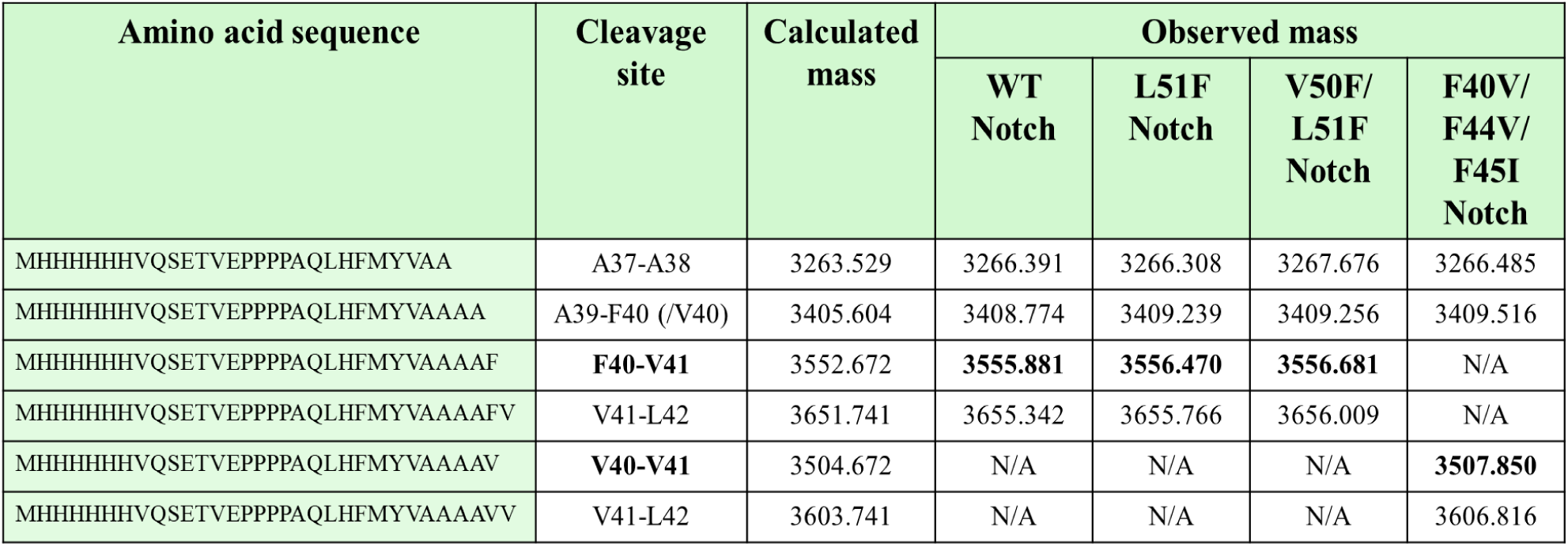
Nβ-like products observed from γ-secretase-mediated cleavage of Notch1 substrate variants. Bold text indicates the major Nβ species. N/A indicates products that cannot be observed due to substrate sequence variation.

These data are consistent with previous reports that the TMD of the substrate is sufficient for interaction and proper cleavage by γ-secretase. After confirming that this double-tagged truncated form of human Notch1 can be used for further studies, we wanted to explore the effects of installing Phe in the P2’ position near the S3 cleavage site in the Notch1 TMD as well as the effect of removing all natural Phe residues in this TMD sequence on the S3 and S4 cleavage products generated by γ-secretase. We, therefore, developed three different Notch1 mutant substrates from the truncated WT Notch1 substrate either by mutating natural amino acids to Phe or by replacing natural Phe residues found in the TMD with the corresponding residue in the APP TMD. All tagged, truncated Notch1 mutants used in this study are identical to the WT Notch substrate except for the mutation sites, as shown in Table 3, and were expressed, purified and validated in a manner equivalent to that of WT Notch. (Fig. S1).

**Table 3.**
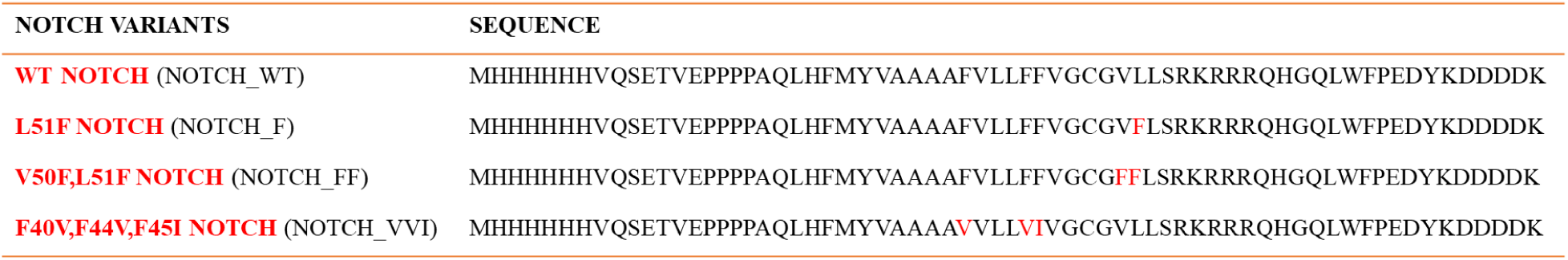
Sequences of Notch variants used in this study aligned with the WT Notch sequence.

### L51F Notch (Notch_F) shifts the S3 cleavage site with reduced product formation

We previously reported that γ-secretase does not tolerate a Phe residue in the P2’ position relative to any cleavage site in APP substrate, resulting in shifting of the cleavage site.^31^ Whether this rule applies to other substrates was not known. To explore this, we designed the L51F Notch1 mutant (Table 3), installing Phe at the P2’ position relative to the S3 cleavage site that generates the NICD-like product.

Purified recombinant L51F Notch1 was incubated with γ-secretase under similar conditions to that of WT Notch1 as previously mentioned. The reaction products were also similarly analyzed by western blotting and MALDI-TOF MS. WT Notch1 substrate was incubated and analyzed in parallel for comparison, and the enzyme reactions with γ-secretase inhibitor were included as controls. As seen in Fig. 3A-D, formation of both NICD- and Nβ-like products were reduced with L51F Notch compared to WT Notch, and no products were seen in any of the control lanes, confirming that the products result from enzyme action.

**Figure 3.**
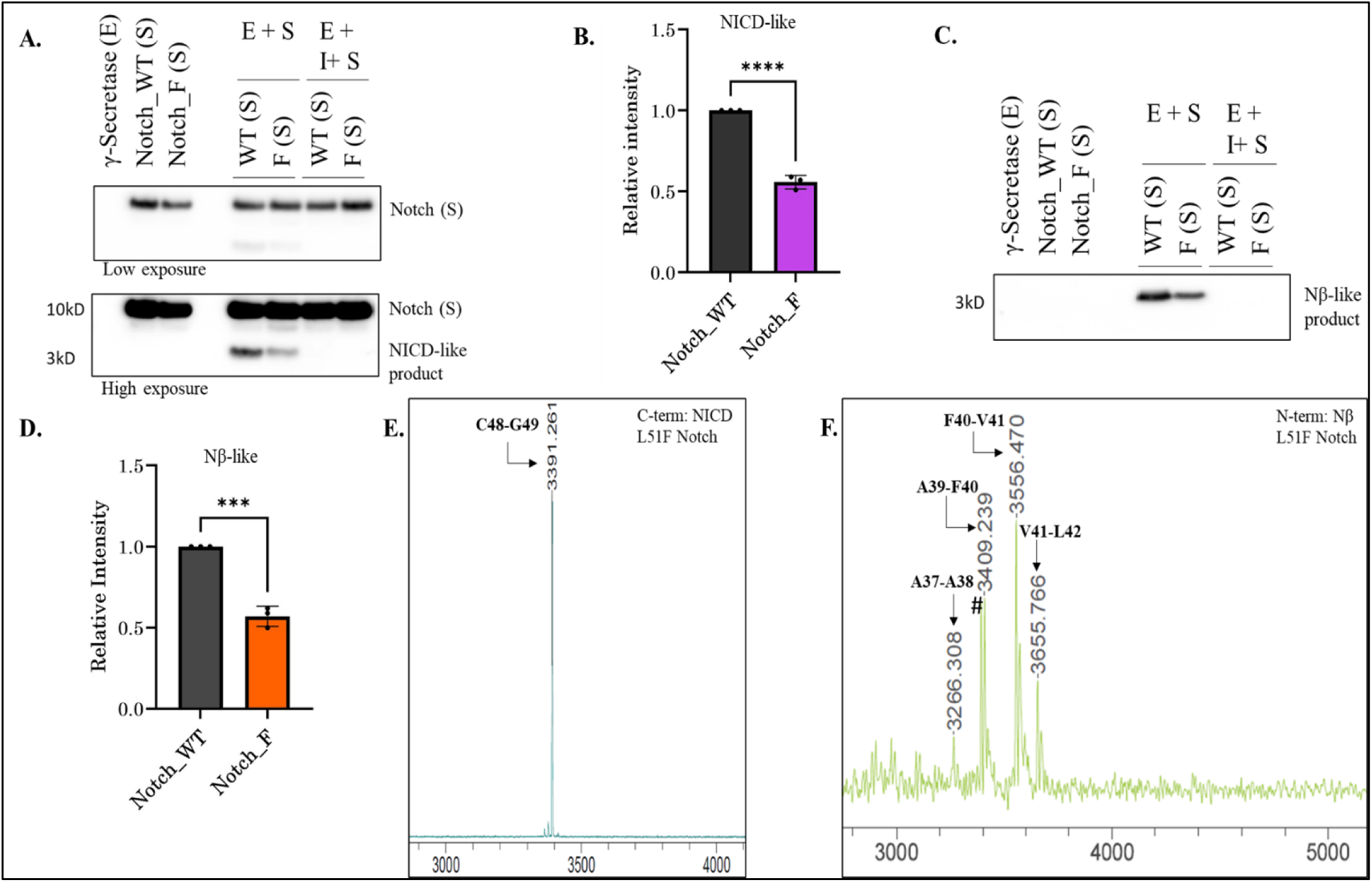
WT vs. L51F Notch1 substrate and analysis of products of γ-secretase processing. A. Immunoblot showing formation of NICD-like products using anti-Flag antibody. NICD-like product formation is reduced in the enzyme reaction with L51F Notch1 mutant compared to reaction with WT Notch1 substrate. B. Densitometry of NICD-like product band from WT Notch1 and L51F Notch1 reactions. C. Immunoblot showing formation of Nβ-like products using anti-His antibody. Nβ-like product formation is reduced in the enzyme reaction with L51F Notch1 mutant compared to WT Notch1 reaction. D. Densitometry of the Nβ-like product band from WT Notch1 and L51F Notch1 reactions. In both blots, no product formation was observed in the control reactions of enzyme alone (E), substrate alone (S), or enzyme + substrate + inhibitor (E+S+I). N=3 for each blot, and unpaired two-tailed student t-test was performed, where * p ≤ .05, ** p ≤ .01, *** p ≤ .001, **** p ≤ .0001. E. MALDI-TOF MS spectra of immunoprecipitated C-terminal NICD-like product from enzyme reaction with L51F Notch1 substrate. A single NICD-like product was observed (GVFLSRKRRRQHGQLWFPEDYKDDDDK); calculated mass: 3390.70, observed mass: 3391.261. F. MALDI-TOF MS spectra for immunoprecipitated N-terminal Nβ-like products. The major product from enzyme reaction with L51F Notch1 was MHHHHHHVQSETVEPPPPAQLHFMYVAAAAF); calculated mass: 3552.672, observed mass: 3556.470. # represents C-terminal NICD-like product non-specifically bound and eluted from beads.

The MALDI-TOF MS spectrum of immunoprecipitated FLAG-tagged NICD-like products from the enzyme reaction with L51F Notch1 clearly showed a shift in the cleavage site by one amino acid towards the N-terminus, with cleavage between C48 and G49 (Fig. 3E) and no detectable product of G49-V50 cleavage. Recently we developed a computational model for activation of γ-secretase for cleavage of WT Notch1 and L51F Notch1 (reported as L36F), that provides structural insights for the shift in cleavage site.^41^ Similar to what was seen with WT Notch, MALDI-TOF MS analysis of immunoprecipitated 6xHis-tagged Nβ-like products for L51F Notch1 showed the major peak corresponding to the fragment generated from F40-V41 cleavage (Fig. 3F and Table 2). In addition to the peak corresponding to F40-V41 cleavage, we observed another major peak from the cleavage between A39 and F40. We also observed other small peaks corresponding to A37-A38 and V41-L42 cleavage. Although all of these Nβ-like cleavage products were observed in both WT Notch1 and L51F Notch1, the intensity of A39-F40 cleavage is much higher than A37-A38 from L51F Notch compared to WT Notch. As with WT Notch1, we did not observe any peak for Nβ-like product generated from direct S3 cleavage (Table S1). Taken together, the data indicates that the intolerance of γ-secretase for Phe in the P2’ position of Notch1 substrate leads to not only shift in the site of S3 cleavage but also reduction in the cleavage of this substrate by the enzyme, likely because of a steric clash between this Phe residue with the relatively small S2’ pocket in the active site of γ-secretase.

### Dysfunctional proteolysis of V50F/L51F Notch (Notch_FF)

APP substrate with Phe in the P2’ position relative to both observed ε sites undergoes dysfunctional and deficient proteolysis by γ-secretase, with generation of very low levels of AICD products.^31^ To explore whether this occurs with Notch1 substrate, we designed the double Phe mutant V50F/L51F Notch, installing two consecutive Phe residues at the positions 50 and 51 in the WT Notch sequence. Thus, we would test if V50F mutation would block the alternative C48-G49 cleavage seen with L51F Notch (Table 3). Purified substrate was incubated with purified enzyme as mentioned earlier, running aforementioned control reactions in parallel. Western blot analysis revealed that this double Phe mutant Notch1 substrate is inefficiently proteolyzed by the enzyme, with reduction in NICD- and Nβ-like product formation compared to WT Notch (Fig. 4A-D). Again, the absence of both NICD- and Nβ-like products in the presence of γ-secretase inhibitor (Fig. 4A,C) indicates that their formation is completely dependent on enzyme action.

**Figure 4.**
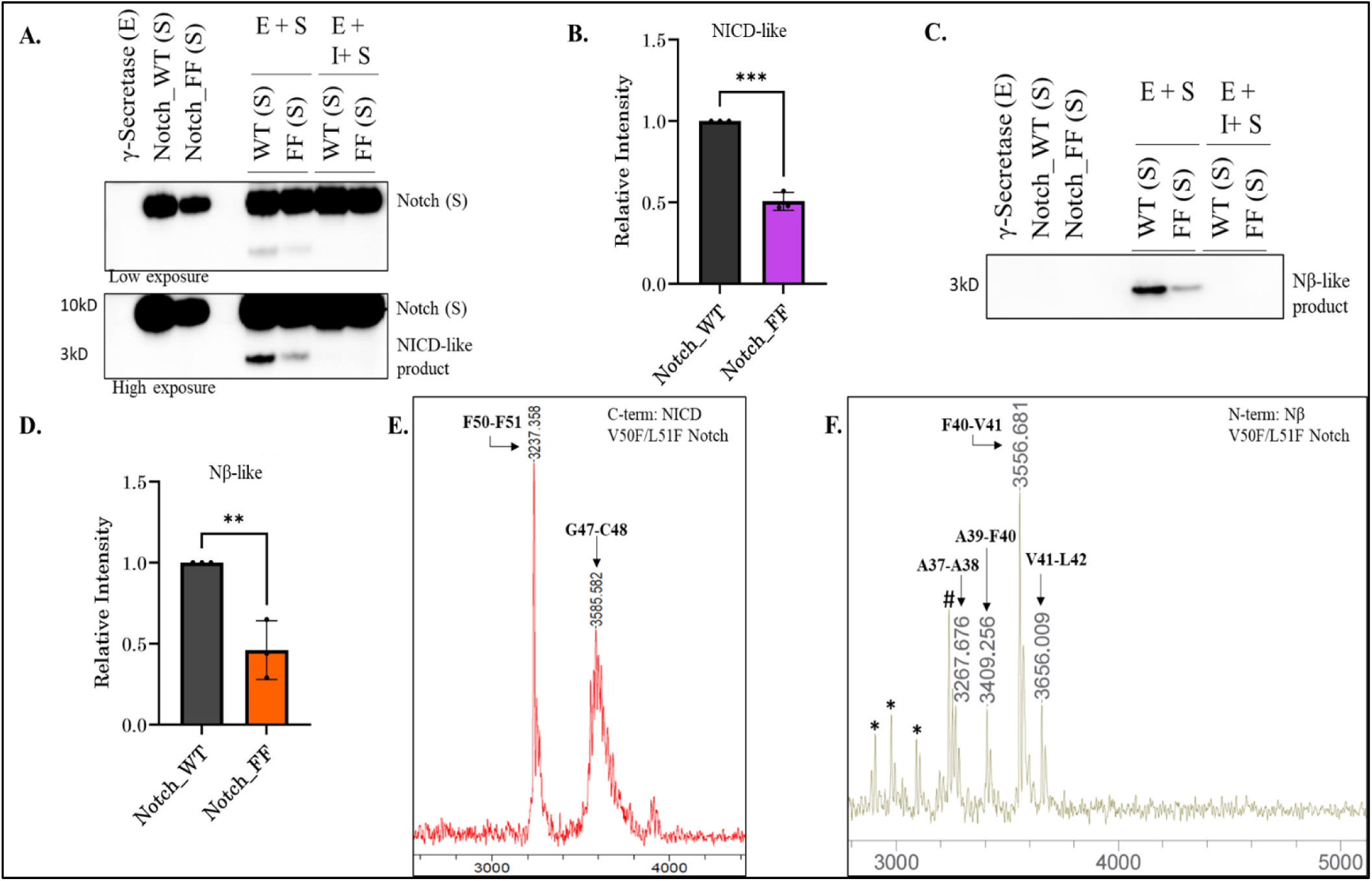
WT vs. V50F/L51F Notch1 substrate and analysis of products of γ-secretase processing. A. Immunoblot showing formation of NICD-like products using anti-Flag antibody. NICD-like product formation is reduced in enzyme reaction with V50F/L51F Notch1 double mutant substrate compared to reaction with WT Notch1. B. Densitometry of the NICD-like product band from enzyme reactions with WT and V50F/L51F Notch1 substrates. C. Immunoblot showing formation of Nβ-like products using anti-His antibody. Nβ-like product formation is reduced in the enzyme reaction with V50F/L51F Notch1 mutant compared to reaction with WT Notch1. D. Densitometry of the Nβ-like product band from enzyme reactions with WT and V50F/L51F Notch1 substrates. In both blots, no product formation was observed in the control reactions of enzyme alone (E), substrate alone (S), or enzyme + substrate + inhibitor (E+S+I). N=3 for each blot, and unpaired two-tailed student t-test was performed, where * *p* ≤ .05, ** *p* ≤ .01, *** *p* ≤ .001, **** *p* ≤ .0001. E. MALDI-TOF MS spectra of immunoprecipitated C-terminal NICD-like products from enzyme reaction with V50F/L51F Notch1 substrate. Two major products were detected: FLSRKRRRQHGQLWFPEDYKDDDDK, with calculated mass of 3234.607 and observed mass of 3237.358, and CGFFLSRKRRRQHGQLWFPEDYKDDDDK, with calculated mass of 3580.669 for the K^+^ adduct and observed mass: 3585.582. F. MALDI-TOF MS spectra for immunoprecipitated N-terminal Nβ-like product from enzyme reaction with V50F/L51F Notch1 substrate. The major product detected was MHHHHHHVQSETVEPPPPAQLHFMYVAAAAF, with calculated mass of 3552.672 and observed mass of 3556.681. * Represents unidentified peak and # represents C-terminal NICD-like product non-specifically bound and eluted from beads.

Interestingly, quantification of the western blots for V50F/L51F Notch showed only about 50% reduction in the NICD-like production compared to WT Notch, in contrast to the sharp reduction in AICD production previously reported with the corresponding double Phe mutant of APP substrate.^31^ Immunoprecipitation and MALDI-TOF MS analysis of the NICD-like products revealed multiple cleavage sites for this substrate. Two major products were seen, resulting from cleavage between F50 and F51 or between G47 and C48 (Fig. 4E, S2), analogous to the low-level products from the corresponding double Phe APP substrate.^31^ As expected, MS did not show any fragments corresponding to cleavage at either G49-V50 (as in WT Notch) or C48-G49 (as in L51F Notch), indicating complete blockage at these sites with V50F/L51F Notch. MALDI-TOF MS analysis of the Nβ-like products showed a major peak resulting from F40-V41 cleavage, in addition to smaller peaks from cleavages at A37-A38, A39-F40 and V41-L42. The S4 cleavage pattern for V50F/L51F Notch is very similar to WT and L51F Notch substrates, with only variations in intensities of the cleavage products. Again, peaks for Nβ products resulting from cleavage at S3 sites were not observed (Fig. 4F and Table S1). Given the multiple NICD-like products observed for S3/ε cleavage and reduced total proteolysis, we conclude that V50F/L51F Notch undergoes dysfunctional proteolysis by γ-secretase.

### Natural Phe residues in Notch1 TMD reduce S3 cleavage

The above results demonstrate that Phe is not tolerated by γ-secretase at P2’ position for S3 cleavage of Notch1, similar to what was seen with ε cleavage of APP substrate. We were then interested in understanding the effects of the three natural Phe residues in the Notch1 TMD. We therefore generated F40V/F44V/F45I Notch1 (Notch_VVI) substrate, by replacing the three natural Phe residues with the corresponding residues from the APP TMD (Table 1 and 3). Western blot analysis for NICD- and Nβ-like products (Fig. 5A-D) showed increased product formation (∼25%) compared to WT Notch. MALDI-TOF MS analysis of the immunoprecipitated products revealed the expected major peak for NICD-like product corresponding to cleavage at the normal S3 G49-V50 site, although very small peaks for products resulting from cleavage at other sites (C48-G49 and G47-C48) were detected as well. Interestingly, the MALDI for Nβ-like products for F40V/F44V/F45I Notch showed the major peak for V40-V41 cleavage and smaller peaks for A37-A38, A39-V40, and V41-L42 (Fig. 5F), similar to S4 cleavages observed for WT, L51F, and V50F/L51F Notch substrates. Again, no Nβ peaks were observed for products from direct S3 cleavages (Table S1). Together, these results show that the natural Phe residues in the Notch1 TMD appear to have little or no effect on the S3 and S4 cleavage sites; however, they do modestly inhibit overall proteolysis by γ-secretase.

**Figure 5.**
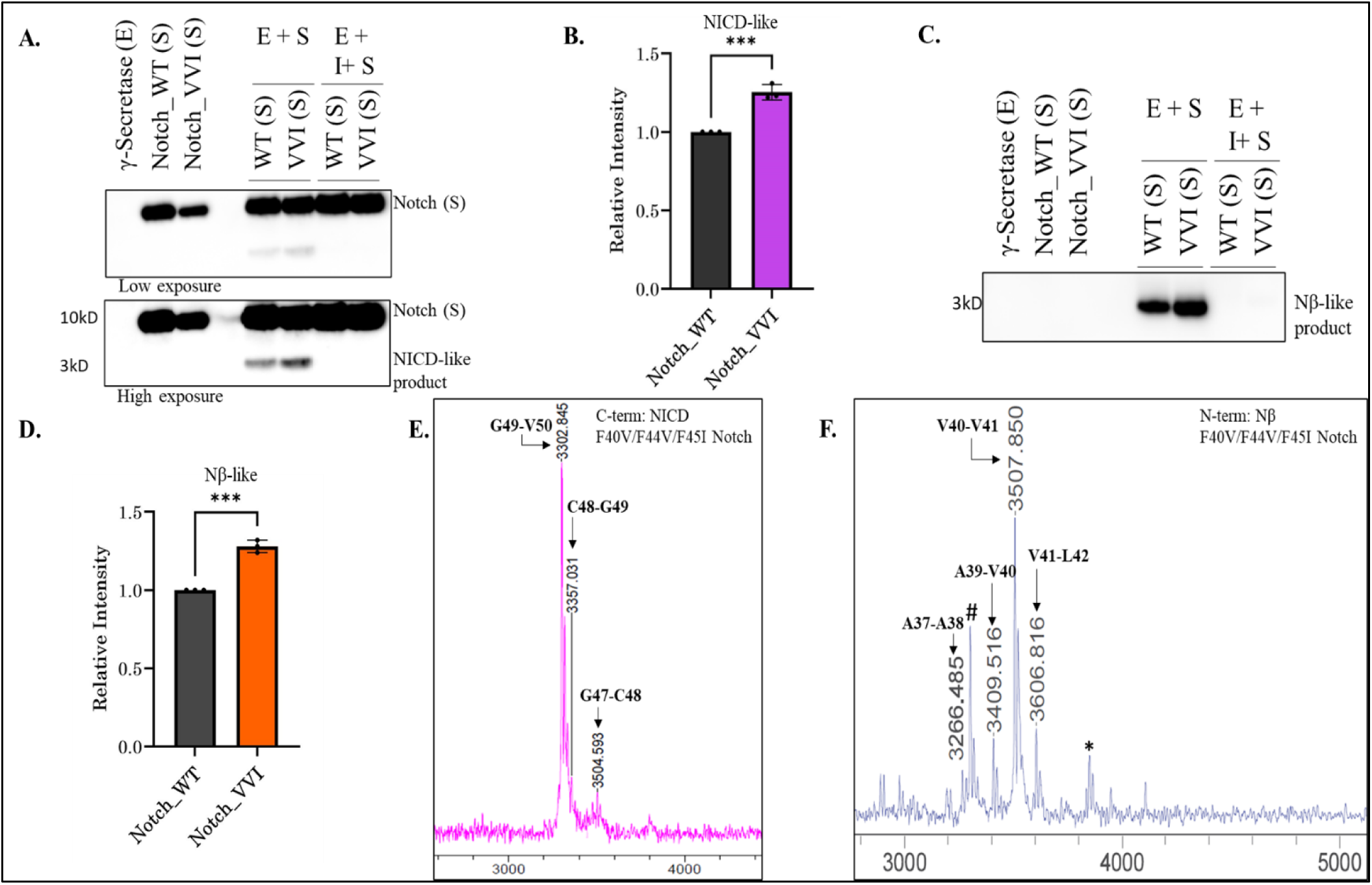
Effects of replacement of three natural Phe residues in the Notch1 TMD on γ-secretase processing. A. Immunoblot showing formation of NICD-like products using anti-Flag antibody from γ-secretase enzyme reactions with WT vs. F40V/F44V/F45I Notch1 substrate. B. Densitometry of NICD-like product bands from the enzyme reactions. C. Immunoblot showing formation of Nβ-like products using anti-His antibody. D. Densitometry of Nβ-like product band from the enzyme reactions. In both the blots, no product formation was observed in the control reactions of enzyme alone (E), substrate alone (S), or enzyme + substrate + inhibitor (E+S+I). N=3 for each blot, and unpaired two-tailed student t-test was performed, where * *p* ≤ .05, ** *p* ≤ .01, *** *p* ≤ .001, **** *p* ≤ .0001. E. MALDI-TOF MS spectra of immunoprecipitated C-terminal NICD-like product from enzyme reaction with F40V/F44V/F45I Notch1 substrate. Products detected include VLLSRKRRRQHGQLWFPEDYKDDDDK (calculated mass: 3299.691; observed mass: 3302.845), GVLLSRKRRRQHGQLWFPEDYKDDDDK (calculated mass: 3356.713; observed mass: 3357.031), and CGVLLSRKRRRQHGQLWFPEDYKDDDDK (calculated mass: 3498.685 for K^+^ adduct; observed mass: 3504.593 (M+K)^+^). F. MALDI-TOF MS spectra of immunoprecipitated N-terminal Nβ-like products from enzyme reaction with F40V/F44V/F45I Notch1 substrate. The major product detected was MHHHHHHVQSETVEPPPPAQLHFMYVAAAAV (calculated mass: 3504.67205; observed mass: 3507.850). * Represents unidentified peak and # represents a C-terminal NICD-like product that non-specifically bound and eluted from the beads.

### Validation of S3 cleavage sites in the Notch1 substrate variants

To further validate the S3 cleavage sites for these Notch variants, we performed western blot analysis of NICD-like fragments from each enzyme reaction using Cleaved Notch (Val1744) antibody. This antibody specifically detects the NICD formed from Notch1 cleavage at the normal S3 site between human G1753 (here G49) and V1754 (here V50) (analogous to murine G1743 and V1744^22^) and does not recognize uncleaved Notch1 or Notch1 cleaved at other sites. A strong NICD-like product band was detected from WT Notch1 substrate, while no band was seen with L51F or V50F/L51F Notch1 substrates (Fig. 6A,B). These findings were consistent with the MS results showing only the WT Notch1 substrate produces NICD-like product cleaved at G49-V50. However, with F40V/F44V/F45I Notch1 (Fig. 6C), we observed a strong signal for the NICD-like product similar to what is seen with WT Notch, confirming that this Notch1 substrate variant, with the three natural TMD Phe residues replaced with corresponding residues in the APP TMD, is indeed cleaved at normal G49-V50 site. However, quantification of the signal intensity of this NICD-like product formed from F40V/F44V/F45I Notch revealed it to be not significantly different from that of product formed from WT Notch (Fig. 6D). This discrepancy with the increased NICD-like band intensity seen with anti-Flag antibody (cf. Fig. 5A), which detects total NICD-like product level, suggests that the cleavages at sites other than G49-V50 in F40V/F44V/F45I Notch contribute to increased formation of NICD-like product. As an additional cross-check of the site of cleavage, we ran a semiquantitative sandwich enzyme-linked immunosorbent assay (ELISA) that detects levels of NICD-like product resulting from cleavage at G49-V50. This ELISA likewise showed no significant difference in signal between enzyme reactions with WT Notch1 and F40V/F44V/F45I Notch1 (Fig. S3). Enzyme reactions with L51F Notch and V50F/L51F Notch showed no signal, as expected, and thereby served as negative controls (Fig. S3).

**Figure 6.**
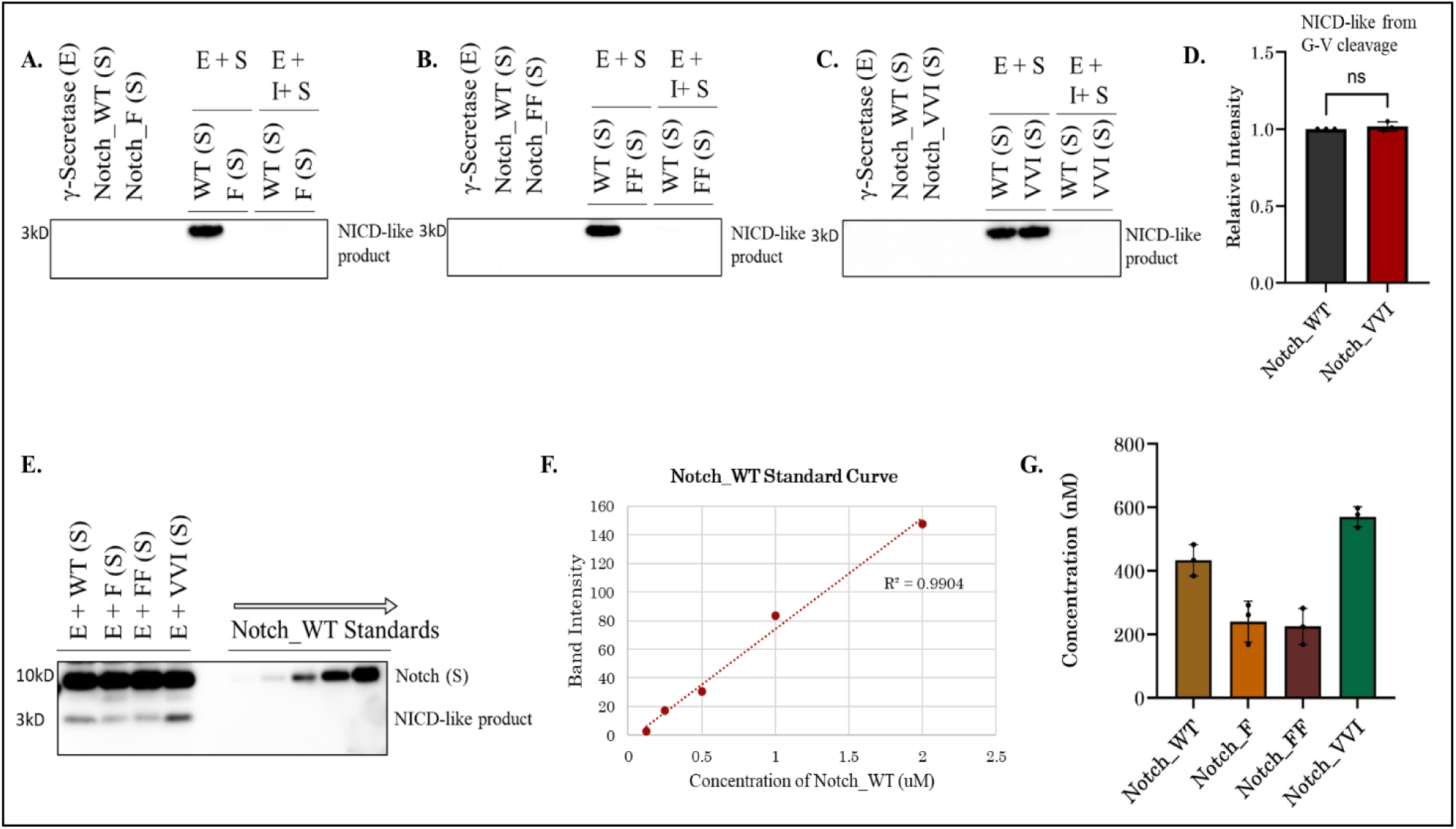
Probing NICD-like products with a neoepitope-specific antibody. Immunoblots for NICD-like products from γ-secretase reaction with each Notch1 substrate. A. Blot from reactions with WT vs. L51F Notch1 substrate using antibody 1744, which specifically recognizes N-terminus of NICD beginning at V50, produced from cleavage at the normal physiological S3/ε cleavage site, at G49-V50. Hence, the blot shows an NICD-like band only in enzyme reaction with WT Notch1 substrate. B. Antibody 1744 blot from reactions with WT vs. V50F/L51F Notch1 substrate. C. Antibody 1744 blot from reactions with WT vs. F40V/F44V/F45I Notch1 substrate. Normal S3/ε cleavage is detected with this mutant substrate. D. Densitometry comparing NICD-like products formed from WT vs. F40V/F44V/F45I Notch1 substrate detected with antibody 1744. Unpaired two-tailed student t-test showed that the difference is nonsignificant (*ns*). E. The western blot for total NICD-like product (probed with anti-Flag antibody) formed from WT, L51F, V50/L51F and F40V/F44V/F45I Notch1 substrates. Run alongside is a known concentration gradient of purified WT Notch1 substrate to generate a standard curve shown in F. for quantitation. G. Calculated concentrations of total NICD-like products formed from each Notch1 substrate variant. N = 3 for all experiments.

To more rigorously quantify levels of total NICD-like products by anti-Flag western blot, we generated a standard curve using known concentrations of purified recombinant WT Notch1 protein alongside the enzyme-substrate reactions for all the variants of Notch1 substrate (Fig. 6E). The standard curve gave an excellent correlation of R^2^=0.9904 (Fig. 6F). The concentrations of NICD-like products from each reaction were then calculated (Fig. 6G), with 434±49 nM from WT Notch1, 241±65 nM from L51F Notch1, 225±57 nM from V50F/L51F Notch1, and 571±31 nM with F40V/F44V/F45I Notch1. These results confirm the ∼50% reduction in NICD-like products with L51F and V50F/L51F Notch1 substrate and ∼25% increase with F40V/F44V/F45I Notch1, which is apparently due to S3 cleavage at sites other than G49-V50.

## DISCUSSION

Despite its biological importance, proteolytic TMD processing of the Notch1 receptor by γ-secretase has not been extensively studied. Here we explored the effects of TMD Phe residues on Notch1 proteolysis by γ-secretase in purified enzyme assays. We find that WT Notch1 TMD cleavage by γ-secretase occurs at two general sites, S3 and S4, generating a single NICD-like product (Fig. 7A and Table 4) and a spectrum of Nβ-like products, consistent with previous studies.^22,26^ Indeed, the specific cleavage sites that lead to the observed products were identical, although we observed major S4 cleavage at F40-V41, instead of A37-A38 as in previous cell-based studies.^26,40^ Importantly, no Nβ or NICD products were observed that would have resulted from Phe in the P2’ position ( likely due to steric clash with the active site S2’ pocket^42^), suggesting that the specificity rule excluding P2’ Phe seen with APP substrate also applies to Notch1.

**Figure 7.**
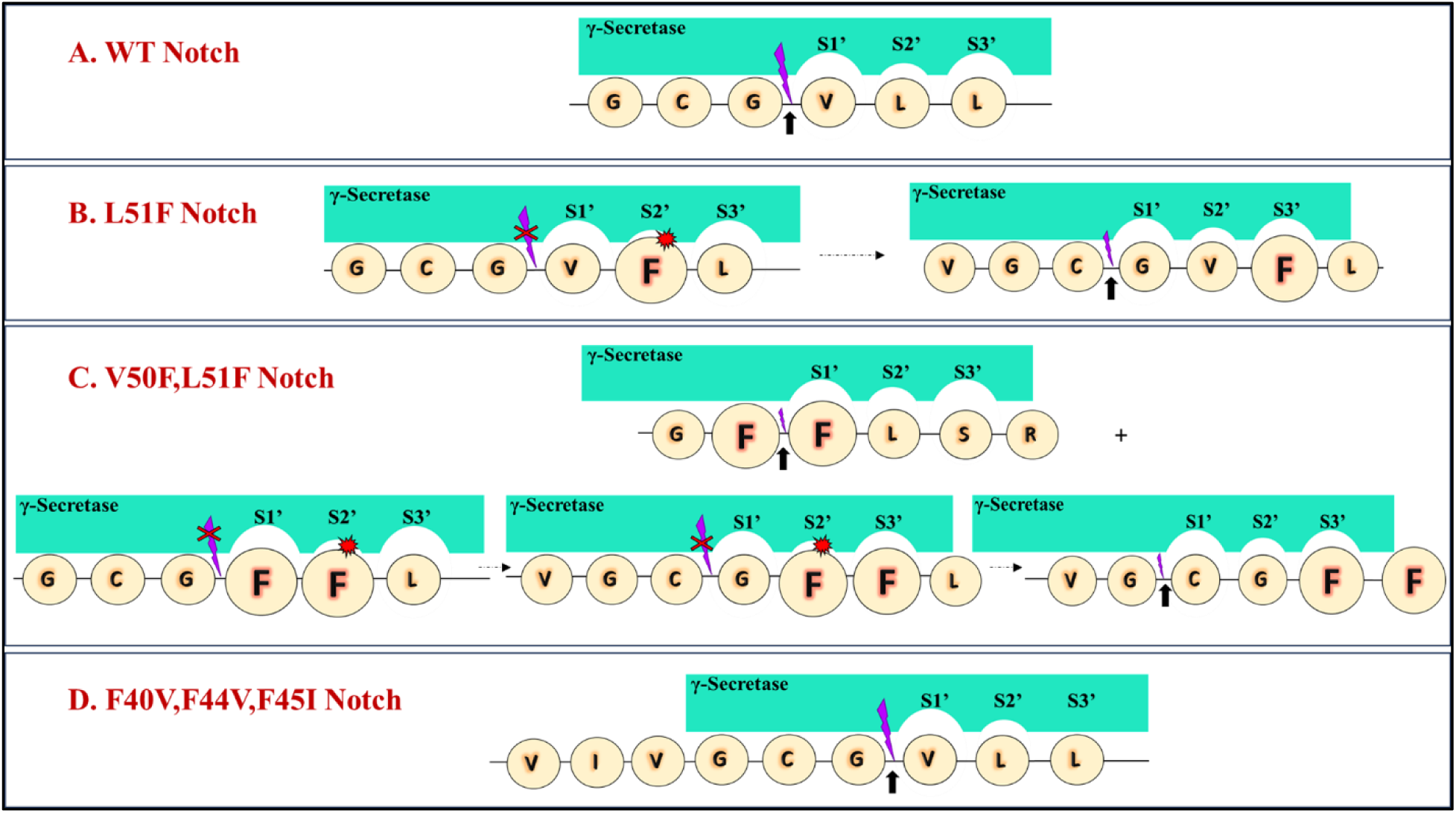
Model for effects of Phe residues on S3/ε cleavage of Notch1 substrate by γ-secretase. Model of the γ-secretase active site showing three active site pockets: S1’, S2’ and S3’. The smaller S2’ pocket does not tolerate Phe in the corresponding P2’ position of substrate. The black arrow points to major observed cleavage sites for each Notch1 substrate variant. (A) WT Notch1 substrate is cleaved at the known G49-V50 site. (B) Cleavage of L51F Notch1 is shifted by one residue, to the C48-G49 site. Cleavage at G49-V50 would place F51 at the P2’ residue, clashing with the small S2’ enzyme pocket. (C) Cleavage of V50F/L51F Notch1 at F50-F51 or G47-C48 prevents placing Phe in the P2’ position for cleavage at either C-48-G49 or G49-F50. (D) Major cleavage of F40V/F44V/F45I occurs at the normal G49-V50 site, as there is no Phe placed in the P2’ position.

**Table 4.**
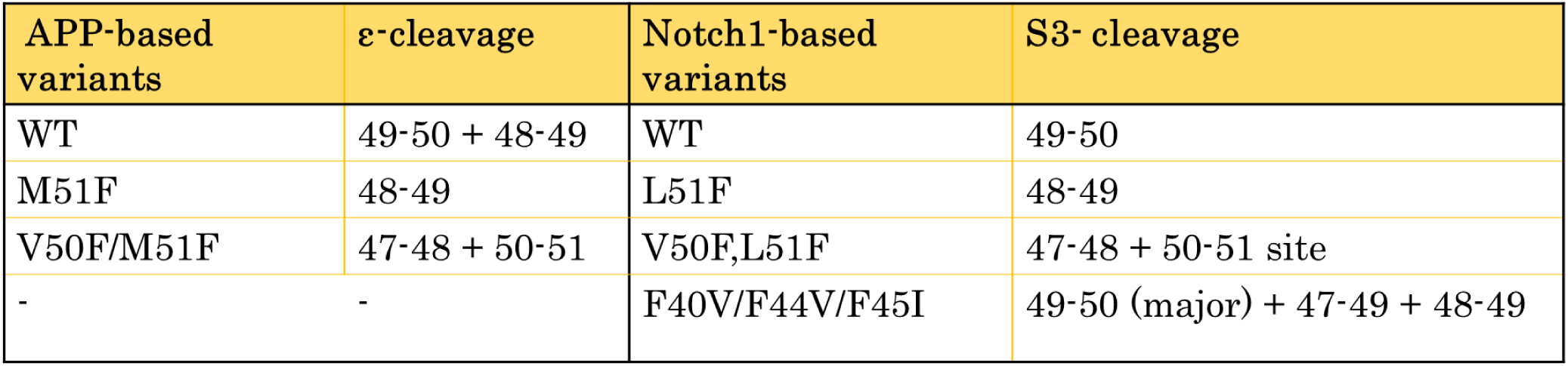
Summary of S3/ε cleavage sites for Notch1-vs. APP-based substrate variants.

To explore whether this P2’ Phe specificity rule indeed applies to Notch1, we first mutated residue L51, which represents the P2’ position for the single S3/ε cleavage site at G49-V50 observed with WT Notch1 substrate. As expected, we observed complete shifting of the S3 cleavage site to C48-G49, with no cleavage between G49-V50, a result consistent with what was seen for the corresponding M51F mutation in APP substrate (Fig. 7B and Table 4).^31^ In contrast, the pattern of Nβ-like products observed with L51F Notch1 substrate was similar to that seen with WT Notch1 substrate. Double Phe mutant V50F/L51F Notch1 substrate blocked S3 cleavage at C48-G49 as well as G49-V50, again consistent with the P2’ Phe specificity rule (Fig. 7C). V50F/L51F Notch1 substrate favors two major S3 cleavages, at G47-C48 as well as F50-F51 sites, again similar to what was seen for the corresponding V50F/M51F mutation in APP substrate (Table 4).^31^ As with L51F Notch1 substrate, the pattern of Nβ-like products observed with V50/L51F Notch1 substrate was similar to that seen with WT Notch1 substrate. This finding is consistent with the regulation of the sites of S4 cleavage by the natural TMD Phe residues present in all three Notch1 substrate variants.

We then explored the role of the three natural Phe residues in the Notch1 TMD. A detailed structure of Notch1 substrate bound to the γ-secretase complex, determined by cryoelectron microscopy^35^, shows these Phe residues all interacting directly with presenilin-1. Thus, they may play a role in S3/ε cleavage by affecting substrate binding and processing. Replacement of all three Phe residues with the corresponding residues in the APP TMD increased proteolysis of the Notch1 substrate. Although the major S3 cleavage site at G49-V50 was maintained (Fig. 7D), two other minor NICD-like products were observed from cleavage at alternative sites (Table 4). Quantification of the major NICD-like product using a neoepitope-specific antibody revealed that the increase in proteolytic processing seen with replacement of the natural Phe residues is apparently due to cleavage at these alternative S3 sites. Thus, the natural Phe residues in the Notch1 TMD are necessary to establish exclusive S3 cleavage at G49-V50, inhibiting extra cleavage events at alternative sites. Interestingly, replacement of the natural Phe residues did not alter the pattern of Nβ-like products observed. The reasons for this are unclear and would require the ability to analyze the small peptide trimming products, as has been done with APP substrate.^9^

Although we show that P2’ Phe residues are not tolerated in Notch1 substrate by γ-secretase, whether S3 cleavage of Notch1 is followed by processive tripeptide trimming is still unknown. We previously provided evidence for three active site pockets (S1’, S2’ and S3’) that dictate tripeptide trimming of initially formed long Aβ intermediates by γ-secretase.^31^ The absence of any long Nβ-like products resulting from direct S3 cleavage in any of the Notch1 substrate variants analyzed strongly suggests that processive trimming, generally in tripeptide increments, could be involved. That may also explain the similar pattern of Nβ-like products observed in all Notch variants tested (Table S2). The presence of multiple Nβ-like species is also consistent with a processive proteolytic mechanism similar to that seen with APP substrate. Whether these processing similarities for APP and Notch1 substrates by γ-secretase extend to other substrates is currently under investigation.

## MATERIALS AND METHODS

### Reagents

Anti-Flag M2 antibody (Sigma #F1804), Cleaved Notch1 (Val1744) antibody (Cell signaling Technology #4147), Anti-His antibody (Invitrogen #MA1-21315), Anti-mouse IgG secondary antibody, HRP (Invitrogen #62-6520), Anti-rabbit IgG HRP linked antibody (Cell signaling Technology #7074).

### Generation of WT Notch1 and mutant constructs

WT Notch construct was designed with His tag on N-terminus and Flag tag on the C-terminus in the pET vector. The WT Notch construct was then mutated using designed primers and Quik Change Lightning mutagenesis kit (Agilent) according to the manufacturer’s instruction to generate Notch mutant constructs L51F Notch, V50F/L51F Notch and F40V/F44V/F45I Notch, and the mutations were verified by sequencing (ACGT, Inc).

### Purification of Notch variants

*E. coli* BL21 cells were transformed with the respective Notch constructs, plated on LB-ampicillin agar and incubated at 37 ^0^C overnight. A single colony was picked for each construct variant and grown with shaking in LB media at 37 ^0^C until OD600 reached 0.8. Cells were induced with 1 mM IPTG and grown for 3 h. Cells were then collected by centrifugation and resuspended in 150 mM NaCl, 10 mM Tris HCl pH 8 and 1% Triton X-100. The cell suspension was passed through a French press, and the lysate was incubated with anti-FLAG M2-agarose beads (Sigma-Aldrich). The beads were then washed with 150 mM NaCl, 10 mM Tris HCl pH 8, 0.25% NP-40. Bound substrates were then eluted from the beads with 100 mM glycine buffer, pH 2.7, with 0.25% NP-40 detergent and stored at −80 °C. The purified proteins were identified and confirmed using MALDI-TOF MS and SDS-PAGE.

### γ-secretase expression and purification

γ-Secretase was expressed in HEK 293F cells by transfection with tetracistronic WT pMLINK vector encoding all four components of the protease complex (gift of Y. Shi, Westlake University^43^). For transfection, HEK 293F cells were grown in suspension in 100 mL of unsupplemented Freestyle 293 media (Life Technologies, 12338-018) until cell density reached 2 × 10^6^ cells/mL. Tetracistronic pMLINK vector (150 ug) was mixed with 25 kD linear polyethyleneimines (PEI; 450 ug) and incubated for 30 min at room temperature. The DNA-PEI mixtures were then added to HEK cells, and cells were grown at 37 ^0^C for 60 h. Cells were harvested, and γ-secretase was purified as described previously.^31^

### Reaction of γ-Secretase and Notch Substrate

All enzyme-substrate reactions were run in the following manner unless otherwise stated. The purified γ-secretase containing presenilin-1 was incubated for 30 min at 37 °C in the assay buffer containing 50 mM 4-(2-hydroxyethyl)-1-piperazineethanesulfonic acid (HEPES, pH 7.0), 150 mM NaCl, 0.1% 1,2-dioleoyl-*sn*-glycero-3-phosphocholine (DOPC), 0.025% 1,2-dioleoyl-*sn*-glycero-3-phosphoethanolamine (DOPE), and 0.25% zwitterionic detergent 3-[(3-cholamidopropyl)dimethylammonio]-2-hydroxy-1-propanesulfonate (CHAPSO), at final enzyme concentration 30 nM. The Notch substrates were then added to final concentrations of 20 µM (for mass spectrometry) or to final concentrations of 10 µM (for western blotting), and the reaction mixtures were incubated at 37 °C for 4 h. The proteolytic products from the enzyme reaction mixtures were then analyzed by mass spectrometry and immunoblotting as described below.

### Detection of NICD-like products by mass spectrometry

NICD-Flag produced from the enzymatic reaction with Notch variants was isolated by immunoprecipitation with anti-Flag M2 beads (Sigma-Aldrich #A2220) in 10 mM MES (2-(4-morpholino) ethanesulfonic acid) pH 6.5, 10 mM NaCl, 0.05% n-Dodecyl-β-D-maltoside (DDM) for 16 h at 4 ^0^C. NICD-like products were eluted from the anti-Flag beads with acetonitrile: water (1:1) with 0.1% trifluoroacetic acid (TFA). The eluants were run on a Bruker autoflex maX MALDI-TOF mass spectrometer.

### Detection of Nβ-like products by mass spectrometry

6xHis-Nβ products produced from enzymatic reactions with Notch variants were isolated by immunoprecipitation with Dynabeads His-tag Isolation & Pulldown (Invitrogen #10103) in 10 mM MES (2-(4-morpholino) ethanesulfonic acid) pH 6.5, 10 mM NaCl, 0.05% DDM detergent for 16 h at 4 ^0^C. Nβ-like products were eluted from the anti-His beads with acetonitrile: water (1:1) with 0.1% trifluoroacetic acid (TFA). The eluants were run on a Bruker autoflex maX MALDI-TOF mass spectrometer.

### Immunoblotting for detection of NICD-like and Nβ-like fragments

Samples from reactions of γ-secretase and Notch variants were subjected to SDS-PAGE on Bis-Tris gels and transferred on PVDF membranes. Membranes were then blocked for 1 h at room temperature in 5% non-fat dry milk in PBST, then incubated with respective primary antibodies for 16 h at 4 ^0^C, followed by washing. The washed membrane was then incubated with corresponding secondary antibodies for 1 h at ambient temperature. Membranes were further washed and imaged by chemiluminescence, and bands were analyzed by densitometry.

### Enzyme Linked Immunosorbent Assay (ELISA)

The reactions of γ-secretase with various Notch substrates (at the final substrate concentration of 5 µM) were performed in the same manner as stated earlier. NICD-like products formed from the reactions that are cleaved only at the known G49-V50 site were detected using PathScan® Cleaved Notch1 (Val1744) Sandwich ELISA Kit from Cell Signaling Technology (#7194) following the manufacturer’s recommended protocol.

### Quantification of NICD-like Fragment

For quantification, WT Notch substrate in assay buffer was used to generate the standard curve in the known concentration range of 0.125-2 µM. The reactions of γ-secretase with various Notch substrates (at the final substrate concentration of 5 µM) were performed in a similar manner as described earlier. Samples were then analyzed by immunoblotting, by probing with anti-Flag M2 antibody.

## AUTHOR INFORMATION

Address: Department of Medicinal Chemistry, School of Pharmacy, The University of Kansas, Lawrence, KS 66045.

## ACKNOWLEDGEMENTS

We thank Dr. Charles Sanders (Vanderbilt University) for providing the Notch1 DNA construct and Dr. Eden Go (University of Kansas Synthetic Chemical Biology Core Laboratory) for technical support on mass spectrometry. This work was supported by Grant AG66986 from the National Institutes of Health to M.S.W.

## SUPPORTING INFORMATION (SI)

MALDI-TOF MS of purified intact Notch1 substrate variants, additional MS spectra for NICD-like product for V50F/L51F Notch1 substrate, ELISA results for NICD-like products resulting from γ-secretase-mediated S3 cleavage at the normal physiological site at G49-V50, tables showing Nβ-like products resulting directly from S3 cleavage and predicted processive proteolysis products of Notch.

