## Supporting Information for "Effects of transmembrane phenylalanine residues on γ-secretase-mediated Notch-1 proteolysis"

**Figure S1. MALDI-TOF MS for purified Notch1-based substrates confirmed the identity for each variant.**

| Notch Variants | Calculated mass | Observed mass |
|---|---|---|
| WT | ююю ы.я ь | ююю ь .юь я |
| Lь шF | юя ч я .я ю | юя чё .шэ э |
| Vь чF/ Lь шF | юя ь э .ё ш | юя ь ю.ё щю |
| Fы ч V/ Fы ы V/ Fы ь I | юэ ы ы. ю ы | юэ ы ы. ы ы ё |

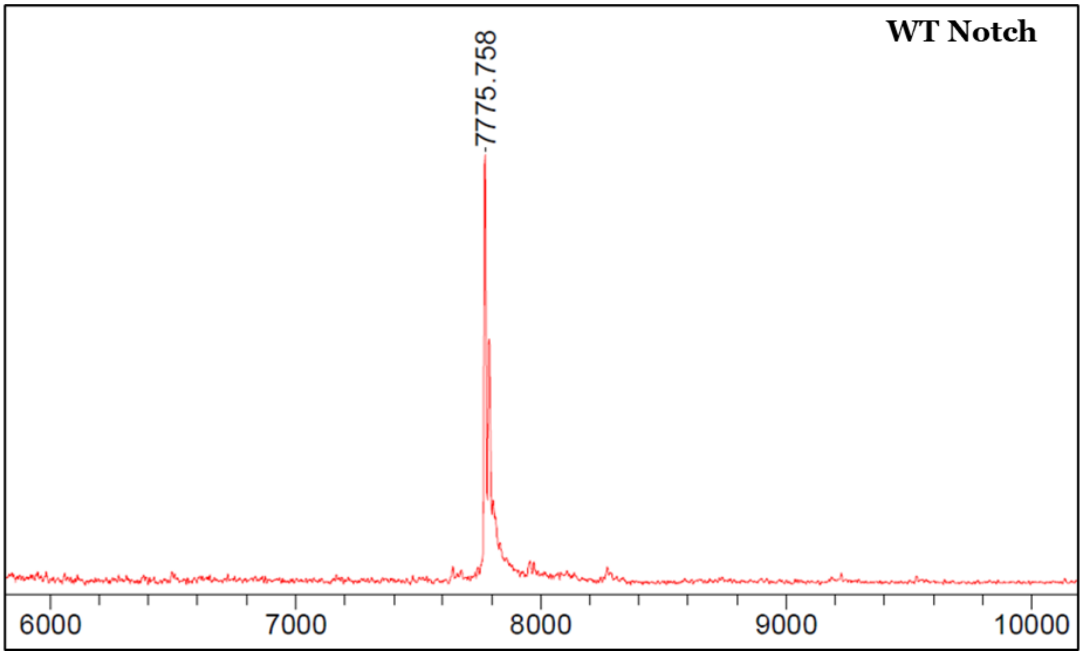

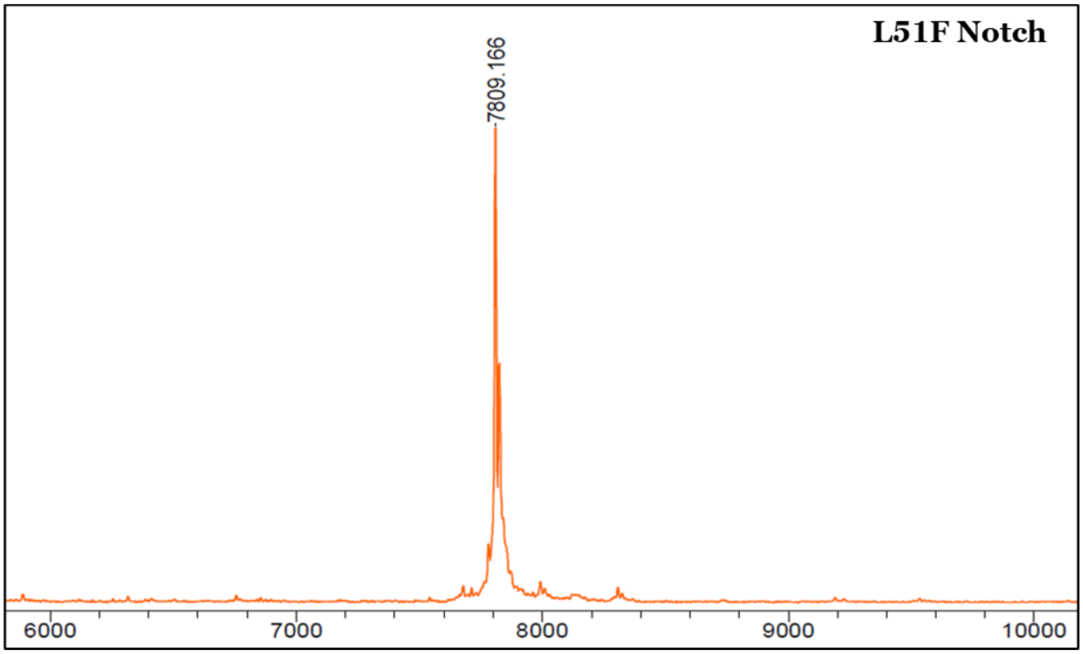
L51F Notch
7809.166
6000
7000
8000
9000
10000

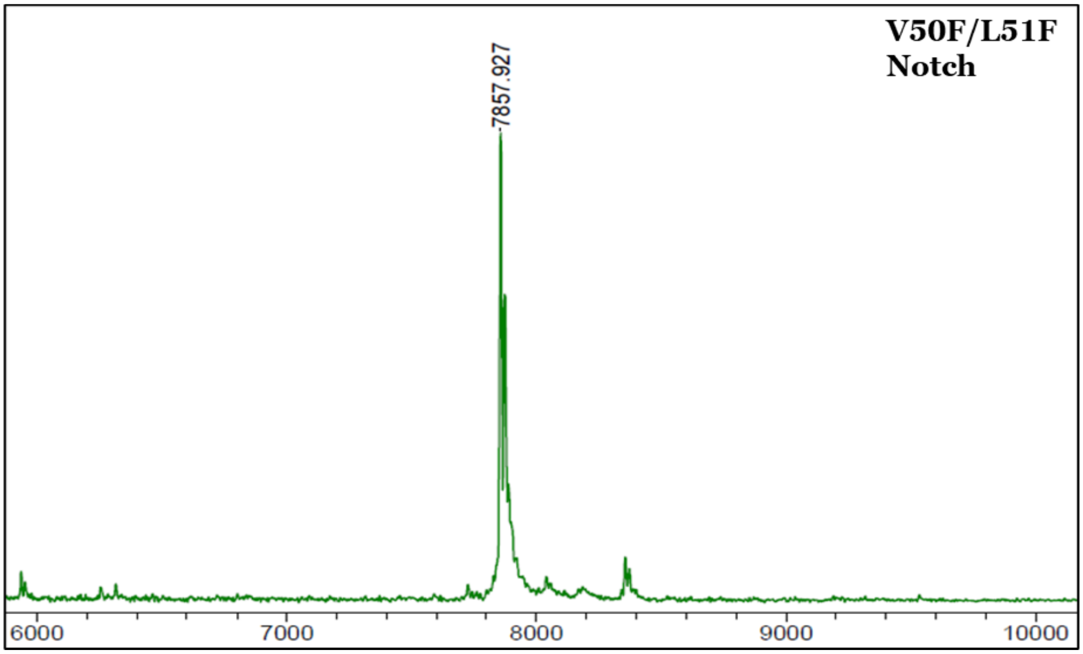
V50F/L51F Notch
7857.927
6000
7000
8000
9000
10000

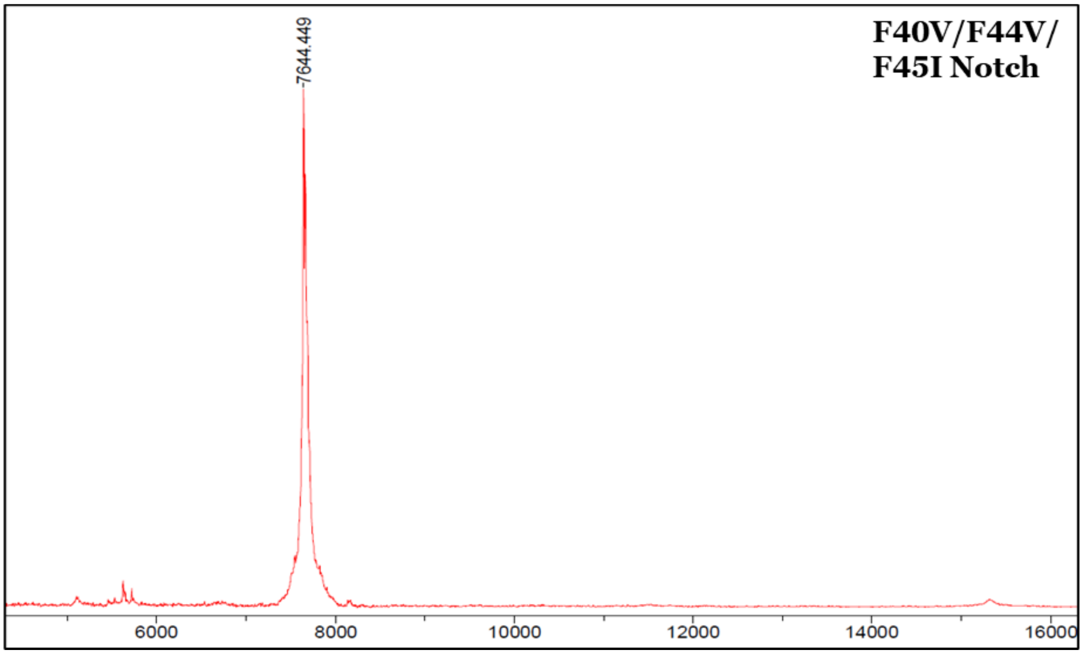

7644.449
F40V/F44V/
F45I Notch
6000
8000
10000
12000
14000
16000

**Figure S2**. **NICD-like product variations for V50F/L51F Notch1 substrate:** C-terminal NICD-like product variations detected by MALDI-TOF MS from γ-secretase reactions with V50F/L51F Notch1 substrate. The same two major products, from cleavage at F50-F51 and G47-C48, were detected in triplicate sample runs, although relative peak intensities varied.

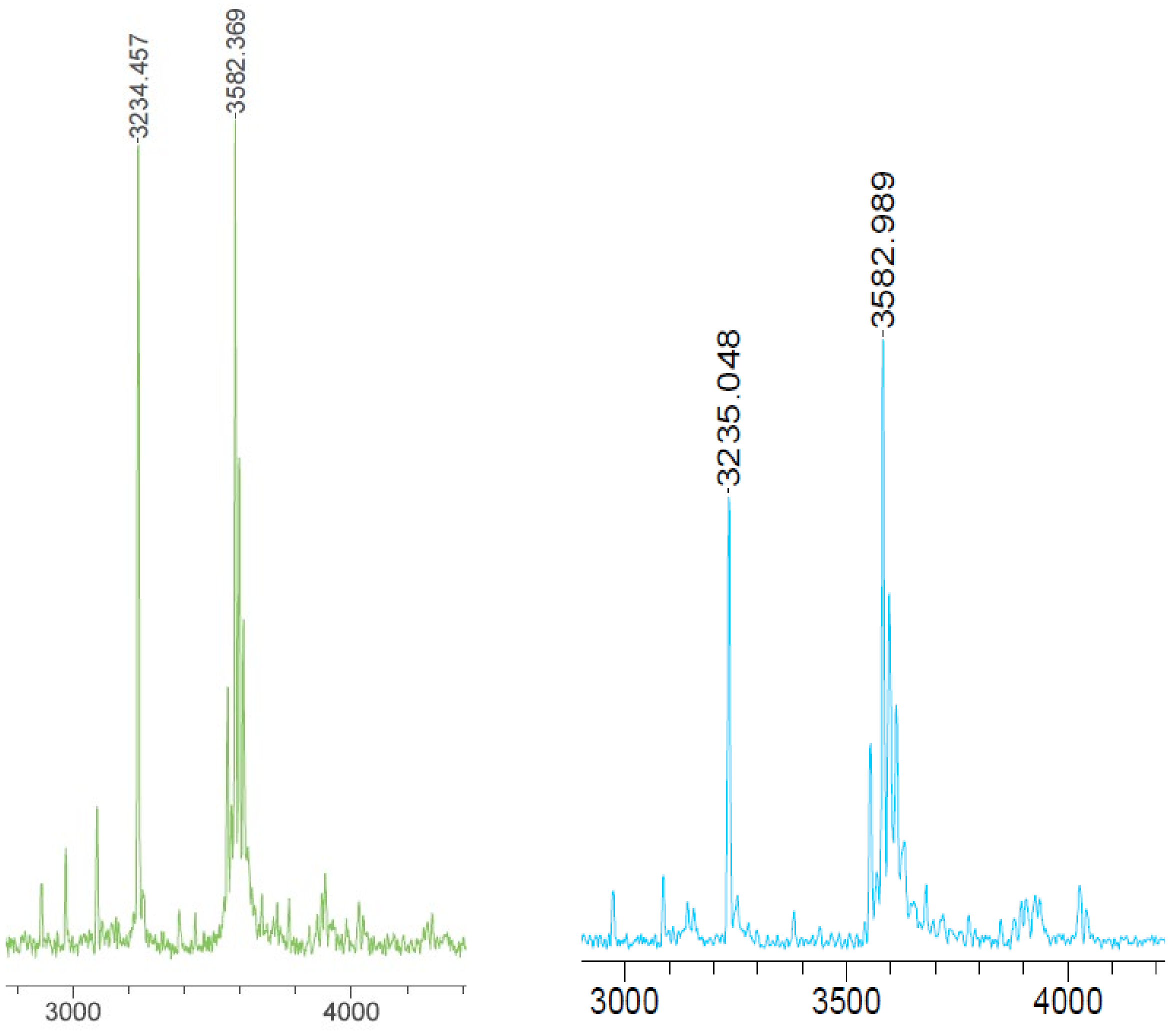

**Figure S3. ELISA results for NICD-like product resulting from γ-secretase-mediated S3 cleavage at the normal physiological site at G49-V50.** Double-antibody ("sandwich") ELISA used Notch1 antibody for capture and a cleaved Notch1 (Val1744) antibody (to neoepitope at N-terminus of NICD) for detection. Optical density was measured at 450 nm (O.D. 450). Note that NICD-like product resulting from γ-secretase cleavage is detected only with WT and F40V/F44V/F45I Notch1 substrates. Unpaired student t-test was performed, n=3.

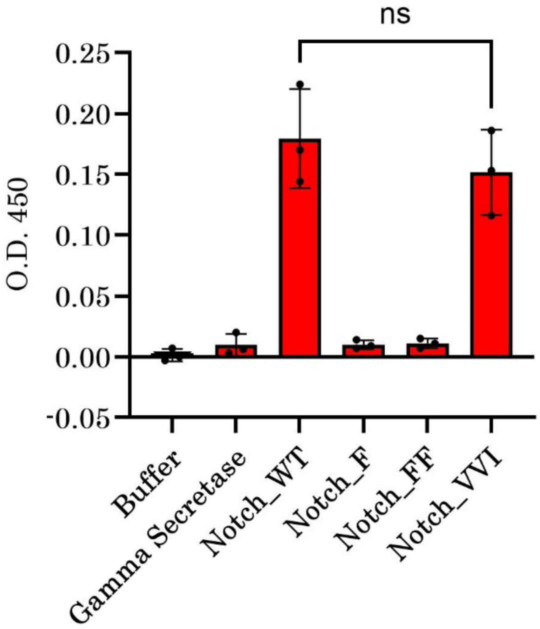

**Table S1. Nβ-like products resulting directly from S3 cleavage.** Sequences of N-terminal Nβ-like products that would directly result from the observed S3 cleavages of the Notch1 substrate variants and their respective calculated masses. None of these products were seen in the MALDI-TOF MS spectra of immunoprecipitated N-terminal products.

| N-term sequence corresponding to the observed S3 cleavages | Calculated mass | Respective Notch variant |
|---|---|---|
| HHHHHHVQSETVEPPPPAQLHFMYVAAAAFVLLFFVGCG | 4357.115 | Notch_WT |
| HHHHHHVQSETVEPPPPAQLHFMYVAAAAFVLLFFVGC | 4300.093 | Notch_F |
| HHHHHHVQSETVEPPPPAQLHFMYVAAAAFVLLFFVG | 4197.084 | Notch_FF |
| HHHHHHVQSETVEPPPPAQLHFMYVAAAAFVLLFFVGCGF | 4504.183 | Notch_FF |
| HHHHHHVQSETVEPPPPAQLHFMYVAAAAVVLLVIVGCG | 4227.130 | Notch_VVI |
| HHHHHHVQSETVEPPPPAQLHFMYVAAAAVVLLVIVGC | 4170.109 | Notch_VVI |
| HHHHHHVQSETVEPPPPAQLHFMYVAAAAVVLLVIVG | 4067.1 | Notch_VVI |

**Table S2. Predicted processive cleavage pattern for Notch variants producing Nβ products observed in MS analysis.** In the table, the various possibilities for cleavage pathways are indicated for WT Notch (rows 1-7). The table also shows only a few representative predicted cleavages for other Notch variants (rows 8-14). Predicted cleavage patterns here avoid placing Phe in P2' position corresponding to any cleavage event and explains observation of similar Nβ spectrum for all Notch variants studied. Notch_VVI would likely undergo cleavage in tripeptide manner shown (as it contains no TMD Phe), yielding similar Nβ products spectrum. After initial cleavage at S3 site, multiple cleavage pathways could be followed resulting in different Nβ-like products observed**.** Amino acids mutations in Notch variants are shown in red.

| No. | Nβ-like product sequence | Predicted small peptides | NICD-like product sequence | Notch variant |
|---|---|---|---|---|
| 1 | MHHHHHHVQSETVEPPPPAQLHFMYVAA | AAFV LLFFV GCG | VLLSRKRRRQHGQLWFPEDYKDDDDK | WT |
| 2 | MHHHHHHVQSETVEPPPPAQLHFMYVAA | AAFV LLF FVGCG | VLLSRKRRRQHGQLWFPEDYKDDDDK | WT |
| 3 | MHHHHHHVQSETVEPPPPAQLHFMYVAA | AAF VLLF FVGCG | VLLSRKRRRQHGQLWFPEDYKDDDDK | WT |
| 4 | MHHHHHHVQSETVEPPPPAQLHFMYVAAAAF | VLLF FVGCG | VLLSRKRRRQHGQLWFPEDYKDDDDK | WT |
| 5 | MHHHHHHVQSETVEPPPPAQLHFMYVAAAA | FVLLF FVGCG | VLLSRKRRRQHGQLWFPEDYKDDDDK | WT |
| 6 | MHHHHHHVQSETVEPPPPAQLHFMYVAAAAFV | LLF FVGCG | VLLSRKRRRQHGQLWFPEDYKDDDDK | WT |
| 7 | MHHHHHHVQSETVEPPPPAQLHFMYVAAAAFV | LLFFV GCG | VLLSRKRRRQHGQLWFPEDYKDDDDK | WT |
| 8 | MHHHHHHVQSETVEPPPPAQLHFMYVAA | AAFV LLFF VGC | GVFLSRKRRRQHGQLWFPEDYKDDDDK | F |
| 9 | MHHHHHHVQSETVEPPPPAQLHFMYVAA | AAFV LLF FVG | CGFFLSRKRRRQHGQLWFPEDYKDDDDK | FF |
| 10 | MHHHHHHVQSETVEPPPPAQLHFMYVAA | AAFV LLF FVG CGF | FLSRKRRRQHGQLWFPEDYKDDDDK | FF |
| 11 | MHHHHHHVQSETVEPPPPAQLHFMYVAA | AAV VLL VIV GCG | VLLSRKRRRQHGQLWFPEDYKDDDDK | VVI |
| 12 | MHHHHHHVQSETVEPPPPAQLHFMYVAAAAV | VLL VIV GCG | VLLSRKRRRQHGQLWFPEDYKDDDDK | VVI |
| 13 | MHHHHHHVQSETVEPPPPAQLHFMYVAAAA | VVL LVI VGC | GVLLSRKRRRQHGQLWFPEDYKDDDDK | VVI |
| 14 | MHHHHHHVQSETVEPPPPAQLHFMYVAAAAVV | LLV IVG | CGVLLSRKRRRQHGQLWFPEDYKDDDDK | VVI |
